# An *In Vivo* Analysis of the Functional Motifs of DEAD-box RNA Helicase Me31B in *Drosophila* Fertility and Germline Development

**DOI:** 10.1101/2022.10.04.510704

**Authors:** Evan Kara, Aidan McCambridge, Megan Proffer, Carol Dilts, Brooke Pumnea, John Eshak, Korey A. Smith, Isaac Fielder, Dominique A. Doyle, Bianca M. Ortega, Yousif Mukatash, Noor Malik, Ammaar R. Mohammed, Deep Govani, Matthew G. Niepielko, Ming Gao

**Author notes:** **Correspondence**, M. Gao, Biology Department, Marram Hall 310, 3400 Broadway, Gary, IN 46408, USA.

## Abstract

In *Drosophila* germline, Me31B is a putative ATP-dependent, RNA helicase that plays role in post-transcriptional RNA regulation to ensure the correct spatial and temporal expression of the mRNAs, a process crucial for proper germline development and fertility. However, Me31B’s *in vivo* working mechanism remains unclear. In this study, we aim to analyze the functions of Me31B’s key domains/motifs to understand how these domains/motifs operate to fulfill the protein’s overall activities. We generated *Drosophila* strains mutant for six important motifs including three ATPase/helicase motifs (DEAD-box, DVLARAK, and HRIGR), the N-terminal domain (N-ter), the C-terminal domain (C-ter), and a protein-binding motif (FDF motif-binding motif). In characterizing these mutants, we observed that the three ATPase/helicase motif mutations cause dominant female sterility which is associated with developmental defects in oogenesis and embryogenesis. Follow-up examination of the DVLARAK motif mutant revealed its abnormalities in germline mRNA localization and transcript level. The Me31B N-ter domain (deletion of C-ter), C-ter domain (deletion of N-ter), and mutation of FDF motif-binding motif led to a decrease in female fertility and abnormal subcellular Me31B localizations in the egg chambers. Moreover, deletion of Me31B N-ter or C-ter motif results in a decrease of Me31B protein levels in the ovaries. This study indicates that these six motifs of Me31B play different roles to contribute to Me31B’s whole-protein functions like ATPase, RNA helicase, protein stability, protein localization, and partner protein binding, which are crucial for germline development and fertility. Considering Me31B protein family’s conserved presence in both *Drosophila* germline and soma (for example, neurons) and in other organisms such as yeast, worm, mouse, and human, the results from this study could expand our understanding of Me31B helicase family’s general working mechanisms in different cell types and species.

## Introduction

*Drosophila* Me31B is an evolutionarily conserved, ATP-dependent, DEAD-box RNA helicase that are important for *Drosophila* germline development and fertility [1–5], with homologs like CGH-1 (*C. elegans*) [6, 7], DjCBC-1 (planarians) [8, 9], Xp54 (*Xenopus*) [10, 11], p54 (mouse) [12, 13], and DDX6/Rck (humans) [14–18] playing similar roles in diverse animal species. In these animals, the main role of Me31B family helicases lies in their post-transcriptional RNA regulation including RNA storage, transportation, translational regulation, stabilization, and decay, which ensure the expression of the messages at the correct time and location, and lead to proper germline functioning and development [14, 19–23].

*Drosophila* Me31B has been employed as a useful model to study the helicases in the family. Me31B’s essentiality for *Drosophila* germline has been underscored by that *me31B* loss-of-function mutation or strong knockdown (KD) cause severe egg chamber development defects or early oogenesis arrest, respectively [1, 24]. In normal egg chambers, Me31B proteins express and aggregate into ribonucleoprotein (RNP) complexes, granular assemblies of proteins and RNAs. In these granules, Me31B complexes with other partner proteins like Tral (a translational repressor protein that usually complexes with Me31B in various types of RNP granules) to render post-transcriptional controls on the RNAs within, a process necessary for proper germline development [3, 25–29]. So far, our understanding of Me31B function mostly came from using *Drosophila* strains with complete loss-of-function alleles of *me31B* gene, strains with a significant loss of Me31B proteins, or biochemical analysis of the protein *in vitro.*Therefore, the protein’s molecular-level working mechanism *in vivo* remains unclear. In this study, we aim to analyze the *in vivo* functions of important domains/motifs of Me31B and then understand how they fit together to contribute to Me31B’s whole-protein activities.

This study focuses on six Me31B domain/motifs (domains and motifs will be called motifs from here for simplicity): DEAD-box motif (AA 207 – 210), DVLARAK motif (AA 97 – 103), HRIGR motif (381 – 385), N-ter motif (AA 1 – 267), C-ter motif (AA 268 – 459), and FDF motif-binding motif (AA 285 – 289). Previous research suggests that the first three motifs (DEAD-box, DVLARAK, and HRIGR) are crucial for the ATPase/helicase activities of Me31B [30–32]; The two large N-ter and C-ter motifs each contain a RecA-like domain and participate in a wide spectrum of activities including ATPase/helicase, protein binding, and assembly into RNP granules [19, 30, 33]; The FDF motif-binding motif enables Me31B to physically bind to the FDF motifs on translational repressor partners like Tral or EDC3 [33–36]. To study these motifs’ *in vivo* functions, we used the CRISPR gene-editing technique and created *Drosophila* strains carrying point mutations that disrupt the motifs’ function or deletion mutations in the *me31B* genes (Figure 1). Analysis of the resulting *me31B* mutants revealed that the six motifs are important for *Drosophila* fertility and germline development, while playing different molecular-level roles in localizing germline RNA, maintaining germline RNA level, stabilizing Me31B protein and localizing the protein, and interactions with Me31B partner proteins.

**Figure 1.**
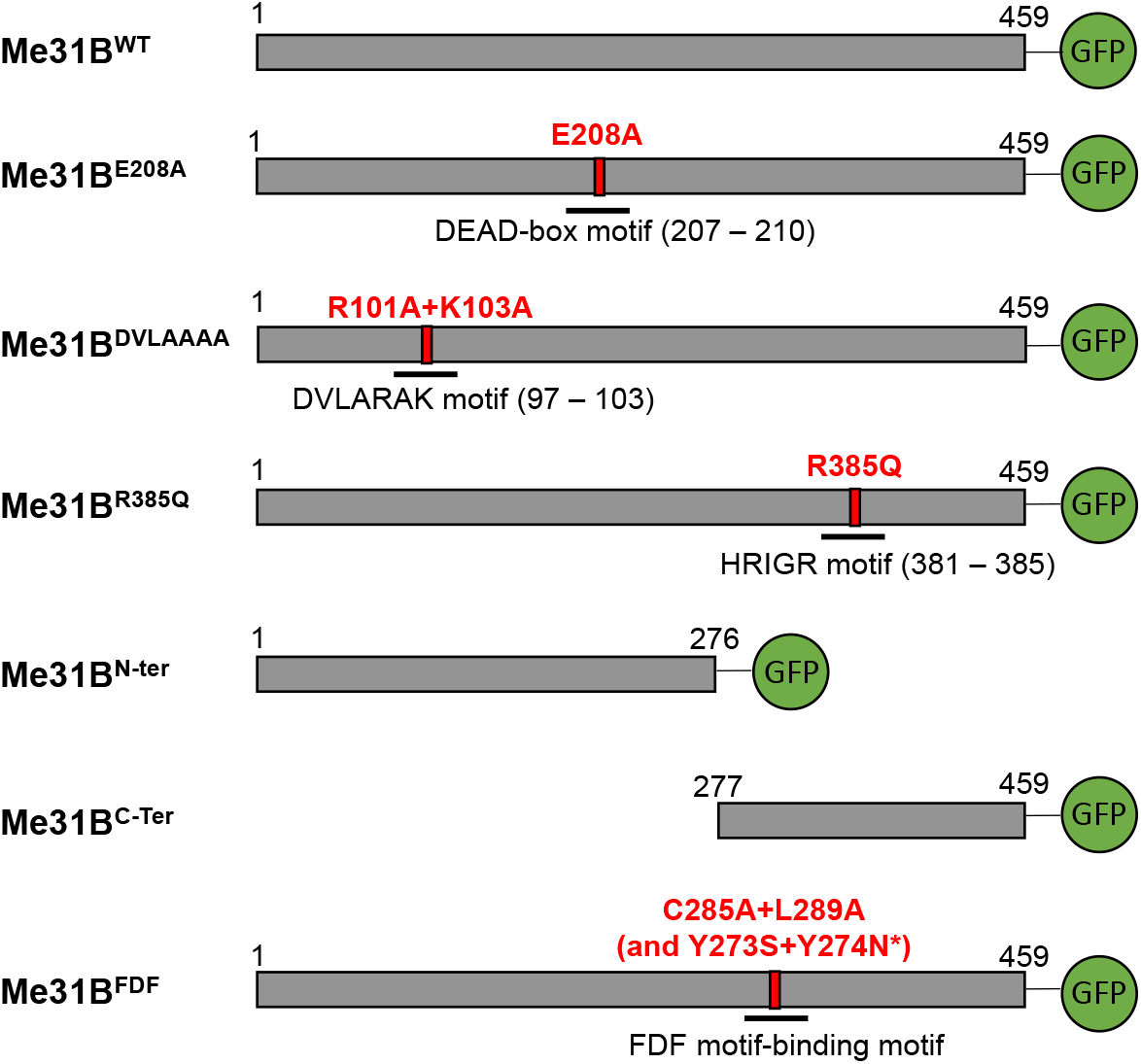
Diagram of the Me31B wildtype and mutant proteins in the *Drosophila* strains generated in this study. The bar shape after each Me31B protein name represents the primary structure of the protein. The numbers on top of the bars mark the number of amino acids and their positions in the protein. Green Fluorescence Protein (GFP) tags are expressed at the C-terminal end of the constructed Me31B proteins. The point mutations are as follows: E208A, glutamic acid 208 replaced by alanine; R101A, arginine 101 replaced by alanine; K103A, lysine 103 replaced by alanine; R385Q, arginine 385 replaced by glutamine; C285A, cysteine 285 replaced by alanine; L289A, leucine 289 replaced by alanine; Y273S, tyrosine 273 replaced by serine; Y274N, tyrosine 274 replaced by asparagine. Me31B^N-ter^ protein (deletion of amino acids 277 to 459) contains the first 276 amino acids of the wildtype protein. Me31B^C-ter^ protein (deletion of amino acids 1 to 276) contains the last 183 amino acids of the wildtype protein. *Note that two unintended, mis-sense mutations were detected in the *me31B^FDF^* strain when sequencing its *me31B* gene. The two mutations (Y273S and Y274N) are outside the FDF motif-binding motif.

## Results and Discussions

### Me31B ATPase/helicase motif mutations lead to dominant female sterility

Although structural and functional studies on Me31B and its homologs identified several ATPase/helicase motifs in Me31B and suggested these motifs’ potential functions in ATP hydrolysis, RNA binding, and RNA translational regulation [11, 19, 30, 32, 34, 37, 38], the *in vivo* functions of the motifs have not been investigated. To study this, we introduced single or multiple point mutations to three conserved ATPase/helicase motifs individually: E208A for DEAD box motif (D**E**AD→D**A**AD), R101A and K103A for DVLARAK motif (DVLA**R**A**K**→DVLA**A**A**A**), and R385Q for HRIGR motif (HRIG**R**→HRIG**Q**). The resulting *me31B* alleles are named *me31B^E208A^, me31B^DVLAAAA^,*and *me31B^R385Q^,* respectively (illustrated in Figure 1). The introduced mutations were reported to disrupt the motifs’ normal functions in Me31B homologs or structurally similar proteins [30–32, 38–41]. When maintaining the three mutant strains, we immediately noticed that all three alleles dominantly cause female sterility. Specifically, *me31B^E208A^/+, me31B^DVLAAAA^/+*, and *me31B^R385Q^/+* heterozygous female flies do not produce any viable progeny in the presence of males from wildtype *w^1118^* strain. In contrast, *me31B^E208A^/+, me31B^DVLAAAA^/+,* and *me31B^R385Q^/+* heterozygous male flies were able to fertilize *w^1118^* females and produce viable progenies bearing the mutant *me31B* alleles, suggesting that the three *me31B* alleles cause sterility in females only.

To find out what caused the sterility, we first checked *me31B^E208A^/+, me31B^DVLAAAA^/+*, and *me31B^R385Q^/+* strains for their embryo laying, development, and morphology. Although embryo laying was observed from all three strains, none (0%) of their embryos (n = 541 for *me31B^E208A^/+*, n = 100 for *me31B^DVLAAAA^/+*, and n = 17 for *me31B^R385Q^/+*)were able to develop to larva or later stages on grape agar plates (see Materials and Methods for the hatchability assay condition). A closer examination of the un-hatched embryos from the three mutants showed that 78% of the embryos from *me31B^E208A^/+*, 33% of the embryos from *me31B^DVLAAAA^/+*, and 100% of the embryos from *me31B^R385Q^/+* had one or more types of patterning defects like fused dorsal appendages, very short, or no dorsal appendages (Figure 2A), consistent with the strains being sterile. Among the three strains, *me31B^R385Q^/+* showed the most severe phenotype: they laid few embryos, and all the embryos laid had severe egg-shell shape defects besides the complete missing of dorsal appendages (Figure 2A).

**Figure 2.**
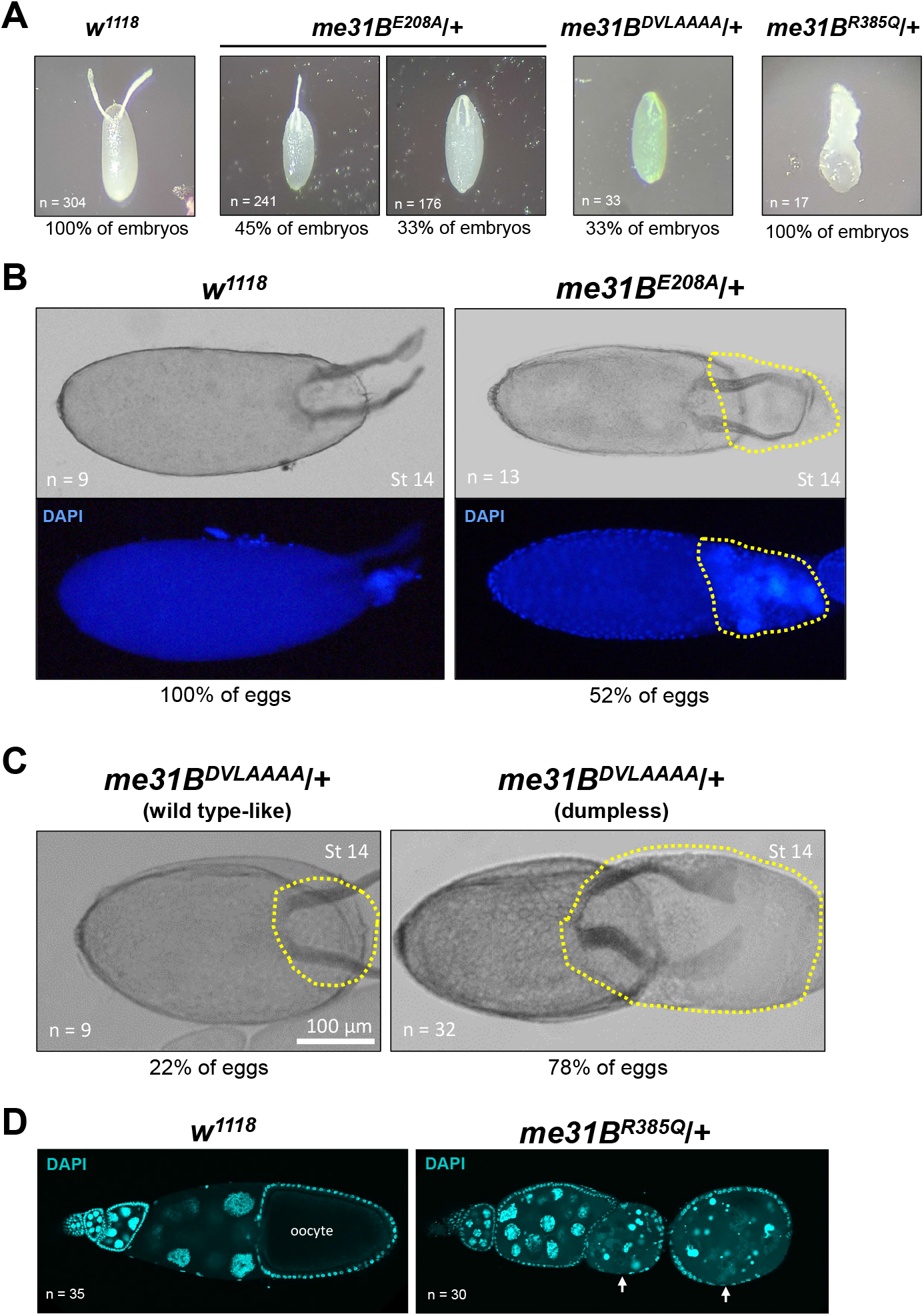
Defective embryogenesis and oogenesis of heterozygous *me31B^E208A^, me31B^DVLAAAA^,* and *me31B^R385Q^* strains. **(A)** Embryos laid by *w^1118^* control females all had normal (100%) dorsal appendages. In contrast, embryos laid by *me31B^E208A^/+*females had fused (45%), shortened (33%), and normal (22%, not shown) dorsal appendages; embryos laid by *me31B^DVLAAAA^*/+ females had shortened (33%) or normal (67%, not shown) dorsal appendages; embryos laid by *me31B^R385Q^*/+ females all had severe morphological defects (100%). **(B)** In 52% of late-stage (stage 14) eggs of *me31B^E208A^*/+ mutant (right panel), dumping of the nurse cell content into the oocyte was incomplete (see highlighted region with yellow dashed lines), in contrast to the complete dumping of *w^1118^* control eggs (left panel). DAPI staining of the same eggs was shown at the bottom panels, and nurse cell nuclei-like materials were present in the “un-dumped” region of the *me31B^E208A^*/+ mutant egg. **(C)** Similar dumpless phenotypes were observed in 78% of *me31B^DVLAAAA^*/+ mutant’s late-stage (stage 14) eggs. The “un-dumped” regions (highlighted with yellow dashed lines) were broader than the *me31B^E208A^*/+ mutant from (B). Only 22% of *me31B^DVLAAAA^*/+ mutant’s eggs appeared normal. **(D)** Ovarioles from *w^1118^* and *me31B^R385Q^*/+ females were stained by DAPI stain. The ovarioles are oriented so that the early-stage egg chambers are on the left and the later-stage egg chambers are on the right. Early-to-mid stage egg chamber degenerations in *me31B^R385Q^*/+ mutant (right panel) are indicated by arrows, in contrast to the successfully developed early- and mid-stage egg chambers in the *w^1118^* control (left panel).

The embryonic defects in the three mutants indicate that their oogenesis did not proceed normally, so we examined their ovaries. In *me31B^E208A^* and *me31B^DVLAAAA^* mutant ovaries, we observed defects in nurse cell dumping, a process in which the nurse cells completely expel all their cytoplasm into late-stage eggs (reviewed in [42]). In the *me31B^E208A^* mutant, 52% of the stage-14 eggs (n = 13) were still connected with egg chamber tissues that contain nurse cell materials (Figure 2B), indicating incomplete nurse cell dumping. This “dumpless” phenotype is more prominent in *me31B^DVLAAAA^* mutant: approximately 78% of *me31B^DVLAAAA^* stage-14 egg chambers (n = 32) did not complete dumping, and the “un-dumped” egg chamber regions were larger than that in the *me31B^E208A^* mutant (Figure 2C). Consistent with this egg development defect, the *me31B^DVLAAAA^* mutant showed reduced egg lay when compared to the wildtype control (Supplementary Figure 1). These phenotypes mimic previously reported dumpless phenotypes in the *dcp-1* (encoding a caspase involved in apoptosis) mutant egg chambers, which also lead to decreased egg lay and early embryo development arrest [43]. The *dcp-1* mutation-induced dumpless phenotypes and sterility were associated with abnormal nucleus envelope breakdown and cytoskeleton organization during nurse cell apoptosis and dumping [43]. Whether the same mechanism is involved in the *me31B^E208A^* and *me31B^DVLAAAA^* mutations remains to be analyzed. The most severe oogenesis defects were observed in the *me31B^R385Q^* mutant, as we observed frequent egg chamber degenerations at early-to-mid stages (Figure 2D), consistent with this mutant’s few egg lay and sterility.

The sterility and defects in oogenesis and embryogenesis in all three mutants suggests that each ATPase/helicase motif is needed for *Drosophila* germline growth, and their cooperative actions are likely needed to output the correct ATPase/helicase function. An explanation for these mutations being dominant could be that the mutations introduced detrimental functions that leads to the mutant Me31B protein being “toxic” to germline cells. This explanation is supported by previous research on similar mutations made in other RNA helicases. For example, in eIF4A, another DEAD-box RNA helicase important for germline mRNA translational control [44, 45], D**E**AD→D**Q**AD and HRIG**R**→HRIG**Q** mutations cause the protein to become dominant negative in translation *in vitro* [46–48]. As another example, in an assay of Xp54 (Me31B homolog in *Xenopus)’s* effect on tethered RNAs, D**E**AD→D**Q**AD mutation, DILA**R**A**K** (homologous to DVLARAK in Me31B)→DILA**A**A**A** mutation, and HRIG**R**→HRIG**Q** mutations changed the protein’s regulation on substrate RNAs from translational repression to translational activation [30]. These reports made us postulate that the dominant sterility effects of *me31B^E208A^, me31B^DVLAAAA^,* and *me31B^R385Q^* mutations may be caused by altered RNA translational control activity of the mutated proteins.

It is integral to note that the HRIG**R**→HRIG**Q** mutation caused much more severe germline phenotypes than the D**E**AD→D**A**AD or DVLA**R**A**K**→DVLA**A**A**A** mutations. We suspect this may be either that 1) substituting an ATPase/helicase motif’s key amino acid (like arginine, R) to glutamine (Q) causes more detrimental effects than alanine (A), or that 2) the HRIGR motif plays a separate and more important roles than the DEAD and DVLARAK motifs. Previous research in other systems provided conceptual support for both explanations. For the first argument, D**Q**AD mutation in the DEAD-box motif of GLH-1 (a conserved germ granule RNA helicase in *C. elegans*) led to a more severe fertility decrease and embryo arrest phenotypes than D**A**AD mutation [49, 50]. For the second, D**E**AD→D**Q**AD mutation in the DEAD-box helicase eIF4A abolish the protein’s ATPase activity but retain a small amount of RNA-binding activity, while the HRIG**R**→HRIG**Q** mutation abolishes RNA-binding ability but retains some ATPase activity [46], suggests that the HRIGR motif may perform unique steps/roles in the helicase’s enzymatic actions.

### Me31B helicase activity functions in *nos* localization by modulating *osk* and *nos* transcript levels

Considering Me31B’s known role in post-transcriptional RNA regulation on important germline mRNAs such as *nanos (nos),* we investigated whether the above ATPase/helicase mutations affect *nos* localization to its germ plasm destination in late-stage eggs (germ plasm is a special cytoplasm at the posterior pole of late-stage eggs and early embryos; germ plasm contains mRNAs needed for processes including embryo patterning and germ cell formation. For reviews, see [51–55]). Normal *nos* localization occurs through the Oskar-dependent formation of homotypic clusters within germ granules, aggregates that contain multiple copies of *nos* transcripts [56, 57]. We tested this hypothesis with the *me31B^DVLAAAA^/+* strain and performed single molecule fluorescent *in situ* hybridization (smFISH) on stage 13 oocytes to identify both unlocalized *nos* transcripts and *nos* clusters in the posterior germ plasm. Using previously established imaging and image analysis techniques [56, 57] (See Material and Methods), we quantified the number of *nos* transcripts that reside within homotypic clusters in the *me31B^DVLAAAA^* mutant and a control CRISPR strain (*me31B^WT^-GFP*) with wildtype *me31B* gene (Figure 3A). We found that the average number of *nos* transcripts found in a homotypic cluster of the control was 6.42 ± 0.47 transcripts per cluster which were not significantly different from the previously published average of 7.58 ±0.42 transcript per *nos* cluster in *yellow white* flies (p = 0.12) [57], suggesting that our CRISPR genome modifications like adding the GFP tag does not affect *nos* localization. However, in the *me31B^DVLAAAA^* mutant, the average number of *nos* transcripts found in a homotypic cluster is 2.33 ± 0.21, a significant reduction when compared to the *me31B^WT^-GFP* control (p < 0.0001). Together, these data suggest that the Me31B helicase activity influences *nos* localization to the posterior germ plasm.

**Figure 3.**
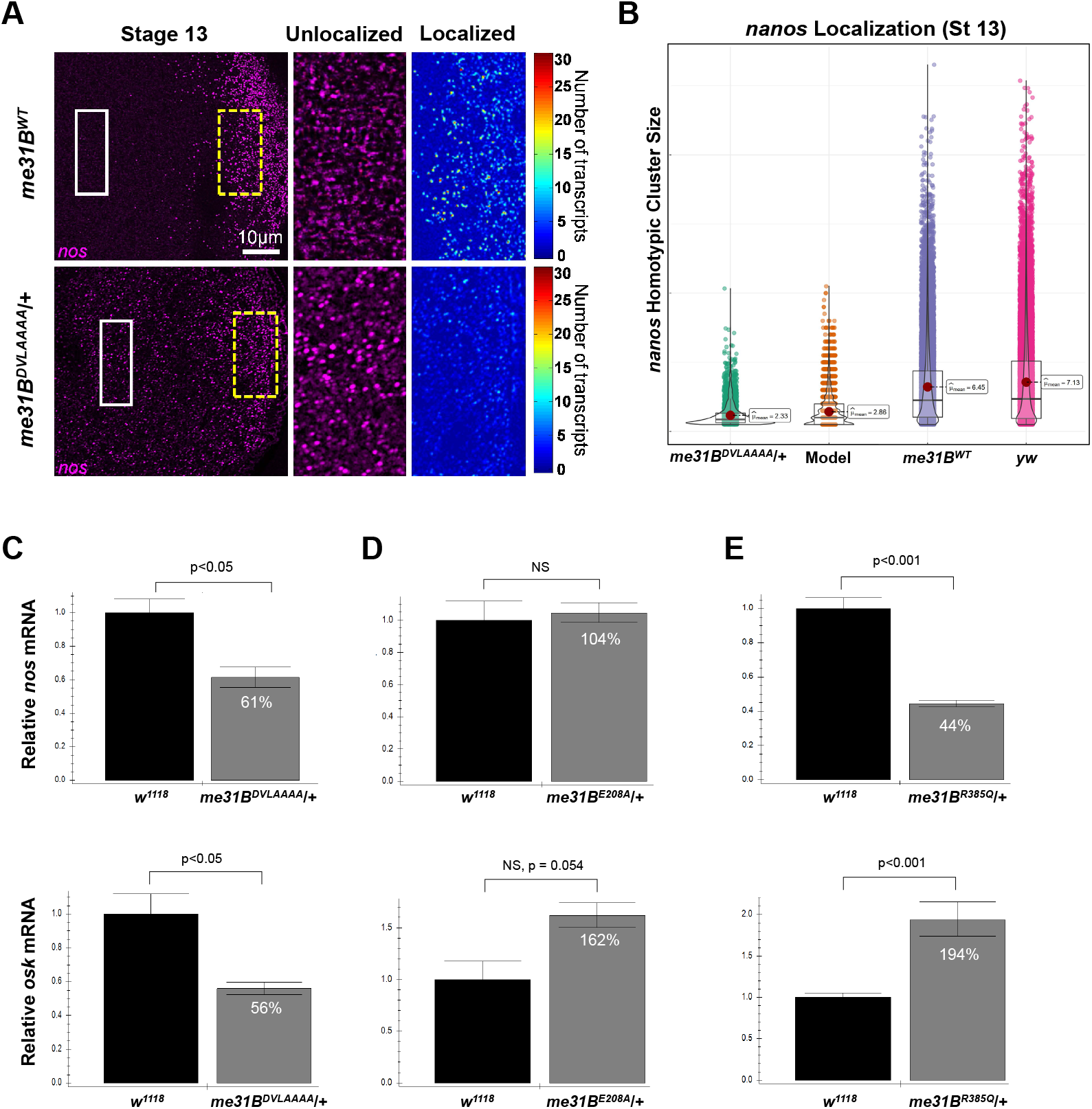
smFISH analysis of *nos* mRNA in *me31B^DVLAAAA^* heterozygous mutant and quantitative RT-PCR analysis of *nos* and *osk* mRNA levels in heterozygous *me31B^E208A^, me31B^DVLAAAA^,* and *me31B^R385Q^* mutants. **(A)** Stage 13 oocytes from *me31B^WT^* and *me31B^DVLAAAA^* flies, with *nos* (magenta) mRNAs detected using smFISH. The white solid box indicates the unlocalized single transcripts of *nos* that are found in the bulk cytoplasm (enlarged in the second panel) while the broken yellow box highlights *nos* that has localized to the germ plasm by forming homotypic clusters (enlarged and shown as a heatmap in the third panel). **(B)** The distribution of *nos* cluster size (number of *nos* transcripts calculated within a granule) found in each genotype’s germ plasm and the computational model. Total clusters identified and analyzed include 3,295 from *me31B^DVLAAA^,* 5,551 from the computational model, 19,195 *me31B^WT^*, and 19,147 from *yellow white* (*yw*). Of note, only oocytes that did not display a dumpless phenotype were included in this analysis. **(C)** In *me31B^DVLAAAA^/+* ovaries, *nos* transcript level is 61% ± 7% of that in the *w^1118^* control (*p* < 0.05) and *osk* transcript level is 56% ± 4% of that in the control (*p* < 0.05). **(D)** In *me31B^E208A^/+* ovaries, *nos* transcript level is 104% ± 7% of that in the *w^1118^* control (NS) and *osk* transcript level is 162% ± 12% of that in the control (NS, *p* = 0.054). (E) In *me31B^R285Q^*/+ ovaries, *nos* transcript level is 44% ± 2% of that in the *w^1118^* control (*p* < 0.001) and *osk* transcript level is 194% ± 12% of that in the control (*p* < 0.001). NS, not significant. Error bar represents standard error of the mean.

We next explored the mechanisms by which Me31B influences *nos* localization. Recently, it has been shown using computational modeling and experimental validation that the localization of *nos* is influenced by 1) the number of *nos* transcripts that are expressed, 2) the amount of Osk in the system, and 3) *nos’s* clustering factor, a quantifiable effect that, in conjunction with transcript and Osk levels, regulates the number of transcripts that can accumulate within a homotypic cluster [57, 58]. To identify whether *me31B^DVLAAAA^* affects one or more of these mechanisms, we first conducted RT-PCR to measure the level of *nos* and *osk* RNAs in the system. We found that *nos* transcript levels were reduced to 61% ± 7% of wild-type levels and *osk* levels were reduced to 56% ± 4% when compared to wild-type (p < 0.05) (Figure 3C). To determine whether these changes can account for the observed reduction of *nos* homotypic cluster sizes in the *me31B^DVLAAAA^* mutant, we used computational modeling to recapitulate *nos* localization *in silico* by adjusting a previously published model’s parameters [58] from 1.00 to 0.61 for *nos* transcript levels and from 1.00 to 0.56 for Osk levels (See Material and methods). Modeled *nos* homotypic clusters contained an average of 2.86 transcripts per cluster which was not significantly different from the average produced by the *me31B^DVLAAAA^* mutant (p = 0.988) (Figure 3B). Together, these data suggest that the mechanism creating the *nos* localization defect in *me31B^DVLAAAA^* mutant is caused by a combined effect generated by the reduction of *nos* and *osk* levels.

Me31B is a known component of an ATP-dependent assembly of a stable, repressed ribonucleoprotein particle (RNP) containing *nos* [3, 22]. In the *me31B^DVLAAAA^* mutant, ATP-dependent helicase activity is presumably disrupted, causing a reduction of *nos* transcripts (Figure 3C). Combining our new data with previous findings that Me31B functions in post-transcriptional mRNA regulation like RNA stability control [1–3, 28, 36], we suggest that Me31B functions in stabilizing unlocalized RNPs containing *nos* and *osk*, allowing for stable mRNAs to reach and become incorporated into the germ plasm. Furthermore, we suggest that disruption of DVLAAAA motif results in a destabilization of mRNAs that have not localized to the germ plasm, resulting in a decrease in *nos* and *osk* that are available to form the germ plasm and ultimately reducing the *nos* localization. Additional germ plasm mRNAs such as *cycB, pgc,* and *gcl* also diffuse to the posterior RNPs containing a single transcript and form homotypic clusters within germ granules [56]. Thus, future studies should explore whether Me31B’s helicase activity is specific to *nos* or has a more global role in stabilizing unlocalized germ plasm mRNAs. To find out whether *nos* and *osk* transcripts level changes also occur in the other two ATPase/helicase mutants (*me31B^E208A^/+* and *me31B^R385Q^*/+), we measured the transcript levels of the two RNAs in the two mutants. To our surprise, in *me31B^E208A^/+* flies, *nos* level remains comparable (104% ± 7%) to *w^1118^* control strain, while *osk* mRNA level is significantly up-regulated (162% ± 12%) (Figure 3D). In *me31B^R385Q^/+* flies, *nos* mRNA showed a significant decrease (44% ± 2%), while *osk* level showed a significant increase (194% ± 12%) when compared to the control (Figure 3E). Comparing the changes of nos and osk between the three ATPase/helicase mutants, (compare Figure 3C, D, and E), we found that the three dominant-sterile mutations caused nonuniform effects on the levels of *nos* and *osk*. This suggests that the three motifs play different roles in maintain germline mRNA levels or they are responsible for maintaining different RNAs. From this, we further suggest that the change of *nos* and *osk* levels cannot be the only cause for the sterility of the three mutants. Further studies are needed to reveal and contrast how different ATPase/helicase motif mutations globally affects germline RNAs.

### Me31B N-ter motif, C-ter motif, and FDF motif-binding motif mutations decrease female fertility

We next analyzed the functions the Me31B N-ter motif (amino acids 1 – 267), C-ter motif (amino acids 268 – 459), and FDF motif-binding motif (amino acids 285 – 289). The N-ter motif contains a RecA-like domain and contributes to the protein’s ATP binding, helicase activity, and P-body assembly ability [19, 30]. The C-ter motif also contains a RecA-like domain and contributes to the protein’s ATP hydrolysis, helicase activity, RNA translational repression, partner-protein binding, and P-body assembly ability [30, 33, 35, 59]. The FDF motif-binding motif allows Me31B to physically interact with FDF motifs in partner RNA repressor proteins like Tral and EDC3 [3, 4, 28, 33, 34, 36]. To study these motifs’ contributions to Me31B whole-protein activities, we generated *me31B^N-ter^, me31B^C-ter^, me31B^FDF^,* and *me31B^WT^* alleles which contain the N-ter coding sequence, the C-ter coding sequence, point mutations in FDF motif-binding motif, and wildtype *me31B* gene (as control), respectively (illustrated in Figure 1). We do note that our attempts to generate the original N-ter (AA 1 - 267) and C-ter (AA 268 - 459) mutations were hindered by technical difficulties, but we were able to generate N-ter (AA 1 - 276) and C-ter (AA 277 - 459) mutations instead, so the latter constructs were used in this study. We first screened the mutants homozygous for the three alleles (*me31B^N-ter^, me31B^C-ter^,* and *me31B^FDF^*) for their fertility by using a previously reported female fertility assay [60]. In this assay, the egg laying and progeny production (hatchability) were separately analyzed to yield a more complete understanding of how the fertility were affected in these mutants. The assay showed that *me31B^N-ter^, me31BC^-ter^,* and *me31B^FDF^* mutant females showed a significant fertility decrease when compared to the *me31B^WT^* control (Figure 4A). Specifically, in the egg laying part, a *me31B^WT^* control female fly laid an average of 117 eggs during the assay period, while a *me31B^N-ter^, me31B^C-ter^,* and *me31B^FDF^* fly laid an average of 68, 29, and 77 eggs, respectively, a significant decrease of 42.9%, 75.2%, and 34.2%, respectively (Figure 4A). In the progeny production part, a *me31B^WT^* control female produced an average of 106 progeny flies, while a *me31B^N-ter^, me31B^C-ter^,* and *me31B^FDF^* fly produced 47, 0, and 75 progenies, a significant decrease of 55.7 % and 100% and an insignificant decrease of 29.2%, respectively (Figure 4B). The obvious absence of progenies from the *me31B^C-ter^* mutant indicated that the *me31B^C-ter^* females were sterile. To validate this, we collected the eggs (n = 97) from *me31B^C-ter^* females (accompanied by *w^1118^* males) on grape-juice agar plates and examined the eggs’ development. We observed that none of the eggs developed to larva or later stages and that all the eggs showed morphological defects including fused dorsal appendages (n = 66, 68% off the eggs) or no dorsal appendages (n = 31, 32% of the eggs) (Figure 4D), consistent with *me31B^C-ter^* females being sterile.

**Figure 4.**
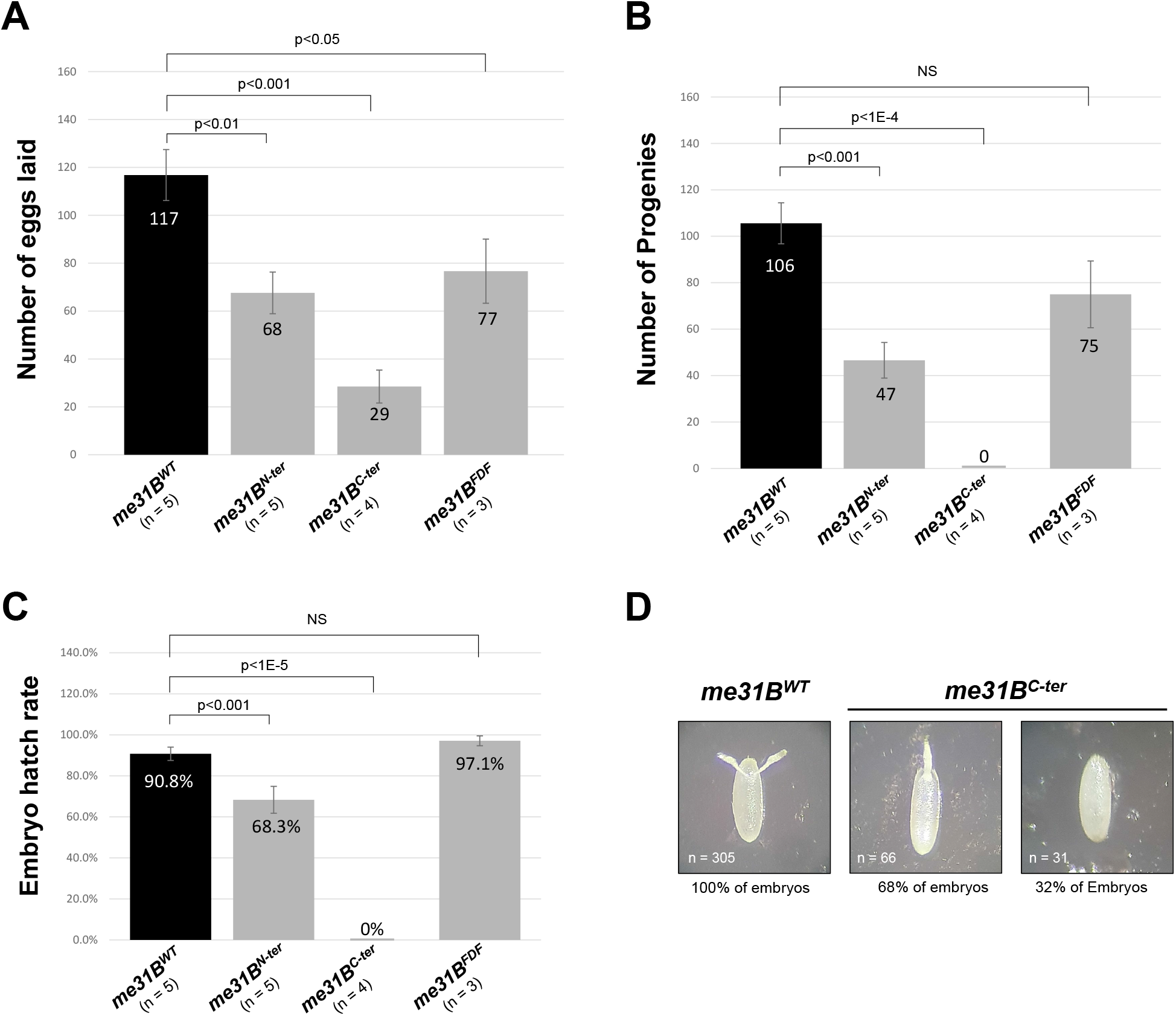
Fertility assays of *me31B^N-ter^, me31B^C-ter^, me31B^FDF^* mutants. **(A)** A *me31B^WT^* (control), *me31B^N-ter^, me31B^C-ter^*, and *me31B^FDF^* female fly laid an average of 117, 68, 29, and 77 eggs, respectively. The egg number of *me31B^N-ter^, me31B^C-ter^*, and *me31B^FDF^* mutants are 58% (*p* < 0.01), 25% (*p*< 0.001), and 66% (*p* < 0.05) of the *me31B^WT^* (control), respectively. **(B)** A *me31B^WT^*(control), *me31B^N-ter^, me31B^C-ter^,* and *me31B^FDF^* female fly produced an average of 106, 47, 0, and 75 progenies, respectively. The progeny number of *me31B^N-ter^, me31B^C-ter^* and *me31B^FDF^* mutants are 44% (*p* < 0.001), 0% (p < 0.0001), and 71% (NS) respectively. NS, not significant. **(C)** The average egg hatch rates for *me31B^WT^* (control), *me31B^N-ter^, me31B^C-ter^*, and *me31B^FDF^* strains are 90.8%, 68.3%, 0%, and 97.1%, respectively. Compared to *me31B^WT^,* the hatch rate decreases in *me31B^N-ter^* and *me31B^C-ter^* mutants are statistically significant but not significant in *me31B^FDF^*. **(D)** Embryos laid by *me31B^C-ter^* flies had fused dorsal appendages (68%) or missing dorsal appendages (32%), in contrast to the normal dorsal appendages (100%) in *me31B^WT^* (control). Error bar represents standard error of the mean.

We calculated the egg hatchability rate of the three mutants by dividing the number of viable progenies by the number of eggs laid. The hatchability rates of *me31B^N-ter^* and *me31B^C-ter^* strains are 68.3% and 0%, respectively, a significant decrease when compared to the control’s 90.8% hatchability rate (Figure 4C). For the *me31B^FDF^* strain, the hatchability (97.1%) is not significantly different from the control (Figure 4C).

We conclude that Me31B’s N-ter or C-ter motifs (when expressed alone) and mutation in the FDF motif-binding motif cause *Drosophila* female fertility decrease. Particularly, *me31B^N-ter^* mutant and *me31B^C-ter^* mutant cause defects in both egg-lay quantity and egg hatchability. However, we cannot neglect that the sterility of *me31B^C-ter^* mutant is a stronger fertility defect than the *me31B^N-ter^* mutant, suggesting the two domains’ different importance in maintaining fertility. Considering that *me31B* is an essential gene and its loss-of-function mutations are lethal [1], we were surprised to see that either Me31B N-ter or C-ter motif (when expressed alone) is enough to support fly viability, suggesting that either motif is able to provide certain essential, full-length-protein functions that enables *Drosophila* growth. Comparatively, *me31B^FDF^* strain showed the smallest fertility decrease with no measurable defects in egg hatchability. This means that the disruption of Me31B’s FDF-motif binding function has only a mild negative effect on fertility.

### Me31B N-ter motif, C-ter motif mutations decrease Me31B protein level

To follow up with the fertility defects in the *me31B^N-ter^, me31B^C-ter^,* and *me31B^FDF^* mutants, we hypothesize that these mutations may have changed Me31B protein stability and therefore reduced the amount of protein in the germline. To test this, we took advantage of the GFP-tags fused to the wildtype and mutant Me31B proteins and used anti-GFP Western blots to quantify the proteins in the ovaries of the mutant strains. We observed that the Me31B^N-ter^ and Me31B^C-ter^ protein levels are 40% and 11% of the Me31B^WT^ control, respectively (Figure 5A and 5B). The Me31B^FDF^ protein level (114%) is comparable to the Me31B^WT^ control. We conclude that the N-ter and C-ter motifs are important for maintaining Me31B protein level and deleting either half of the protein likely destabilized the protein. We also suggest that the Me31B^C-ter^ protein (N-terminal deletion) is even less stable than the Me31B^N-ter^ protein (C-terminal deletion). Together with the fertility experiments (Figure 4), we noticed that the mutant strains’ fertility and their mutant protein expression level share a similar trend, with *me31B^C-ter^* mutant showing complete sterility and the least level of protein expression and *me31B^FDF^* mutant showing a mild fertility decrease and near wildtype-level protein expression (compare Figure 4 and Figure 5B). We speculate that, besides the likely altered Me31B protein functions, the change of protein abundance could also be a factor that decreased the fertility of the mutants.

**Figure 5.**
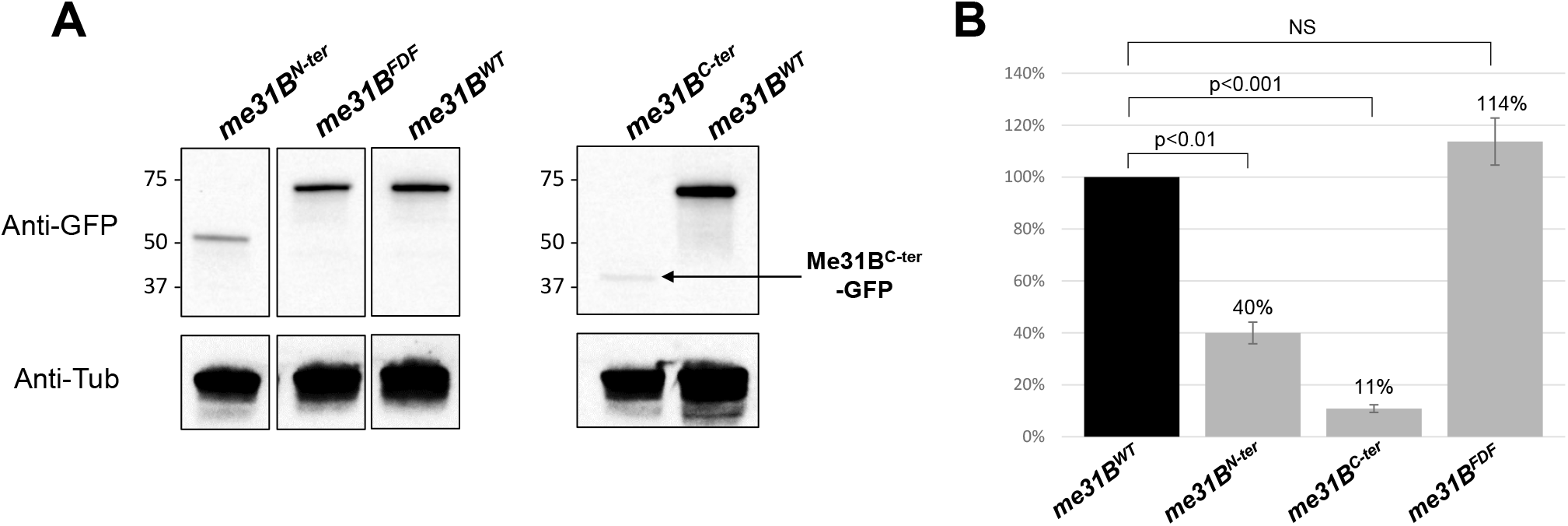
Western blot quantification of Me31B protein level in *me31B^N-ter^, me31B^C-ter^, me31B^FDF^* mutants. **(A)** Anti-GFP Western blots were used to quantify the Me31BWT-GFP, Me31B^N-ter^-GFP, Me31B^C-ter^-GFP, and Me31B^FDF^-GFP proteins in the ovaries of the corresponding fly strains. Anti-Tubulin Western blots were used as loading controls. The expression level of Me31B^N-ter^ and Me31B^C-ter^ proteins were conspicuously lower than the Me31BWT control protein, while the Me31B^FDF^ protein level is comparable to the control. The images shown are representative images of multiple biological replicates. The additional, uncropped biological replicate images are presented in Supplementary Figure 2. **(B)** The Me31B^N-ter^-GFP and Me31B^C-ter^-GFP protein levels are 40% (*p* < 0.01) and 11% (*p* < 0.001) relative to the Me31B^WT^-GFP control protein, respectively. Me31B^FDF^-GFP protein level is 114% relative to the control (NS). Western blot image analysis was performed with ImageJ and protein quantifications were normalized by using the alpha-tubulin proteins. NS, not statistically significant. Error bar represents standard error of the mean.

### Me31B N-ter motif, C-ter motif, and FDF motif-binding motif mutations alter Me31B subcellular localization

Me31B is abundantly expressed in the cytoplasm of nurse cells and developing oocytes in *Drosophila* ovaries. In these cells, Me31B complexes with partner protein Tral and aggregate together into RNP granules including perinuclear granules (nuage granules) in nurse cells, P-body/sponge body granules in nurse cells and oocytes, and germ plasm granules in oocytes [1, 24, 28]. To find out whether the *me31B^N-ter^, me31B^C-ter^,*and *me31B^FDF^* mutations affect Me31B subcellular localization, we used the GFP-tags on the Me31B proteins to visualize them in the ovary cells and used anti-Tral immunostaining to mark the germline RNP granules.

In the early-stage egg chambers (Figure 6A), wildtype Me31B^WT^-GFP and Tral both localize to the nuage granules and the P body/sponge body granules in the nurse cells and developing oocytes (Figure 6A, first row). The two proteins showed extensive overlap at these granules, suggesting likely colocalization. This localization pattern of Me31B^WT^-GFP and Tral is indistinguishable from the wildtype proteins in Oregon-R strains or GFP-Trap strains carrying wildtype *me31B* genes [1, 24]. However, in *me31B^N-ter^, me31B^C-ter^, me31B^FDF^* egg chambers, the three mutant Me31B-GFP proteins were conspicuously more diffused in the nurse cells and oocytes, and they no longer overlap with Tral whose localization remain similar to that in the *me31B^WT^* control (Figure 6A, 3 bottom rows). This indicates that Me31B N-ter motif, C-ter motif, and the FDF motif-binding motif are needed for Me31B’s aggregation status. Furthermore, the three mutant proteins each showed distinct localization/distribution patterns. Me31B^N-ter^-GFP proteins are found in the nurse cell nucleus (Figure 6A, second row), which is further confirmed by co-staining the egg chambers with DNA stain DAPI (Supplementary Figure 3). Although a small number of Me31B^C-ter^-GFP aggregates were observed (Figure 6A, third row), they are fewer and smaller than Me31B^WT^-GFP aggregates. From these observations, we conclude that Me31B’s proper aggregation into germline RNP granules requires the presence of both the N-ter motif and C-ter motif. Previous studies suggest that the two motifs contain different sequence components and therefore different potential functions. The N-ter motif contains several ATPase/helicase motifs including DEAD-box motif and DVLAAAA motif but no known protein-binding motifs [30–32, 35]; The C-ter motif contains several protein-binding motifs and one putative helicase motif, HRIGR) [10, 19, 30, 33, 35, 59]. This means that Me31B likely uses it RNA-interaction ability on both N-ter and C-ter motifs and the protein-binding ability on the C-ter motif to aggregate into the RNP granules. In line with this speculation, the failure of Me31B^FDF^-GFP to aggregate (Figure 6A, bottom row) suggests that interaction/recruitment by FDF-motif proteins like Tral and EDC3 [34, 61] is necessary for Me31B’s aggregation process. We note that Me31B^FDF^-GFP’s failure to aggregate is independent of its protein expression level (Figure 5). Together with the mild fertility decrease of the *me31B^FDF^* strain (Figure 4), we suggest that Me31B’s aggregation status per se only has a small influence on *Drosophila* fertility. Another aggregation factor in Me31B protein is the two predicted Intrinsically Disordered Regions (IDRs) at the N-ter end (AA 1-53) and the C-ter ends (AA 431 – 459), respectively. Deleting the IDRs caused rapid self-assembly of Me31B into aggregate-like structures *in vitro,* so the IDRs were suggested to attenuate the interactions between the folded N-ter and C-ter motifs [62]. From the above pieces of evidence, Me31B aggregation to germline RNP granule could be a complex interplay between RNA-interacting, partner protein binding, and IDRs.

**Figure 6.**
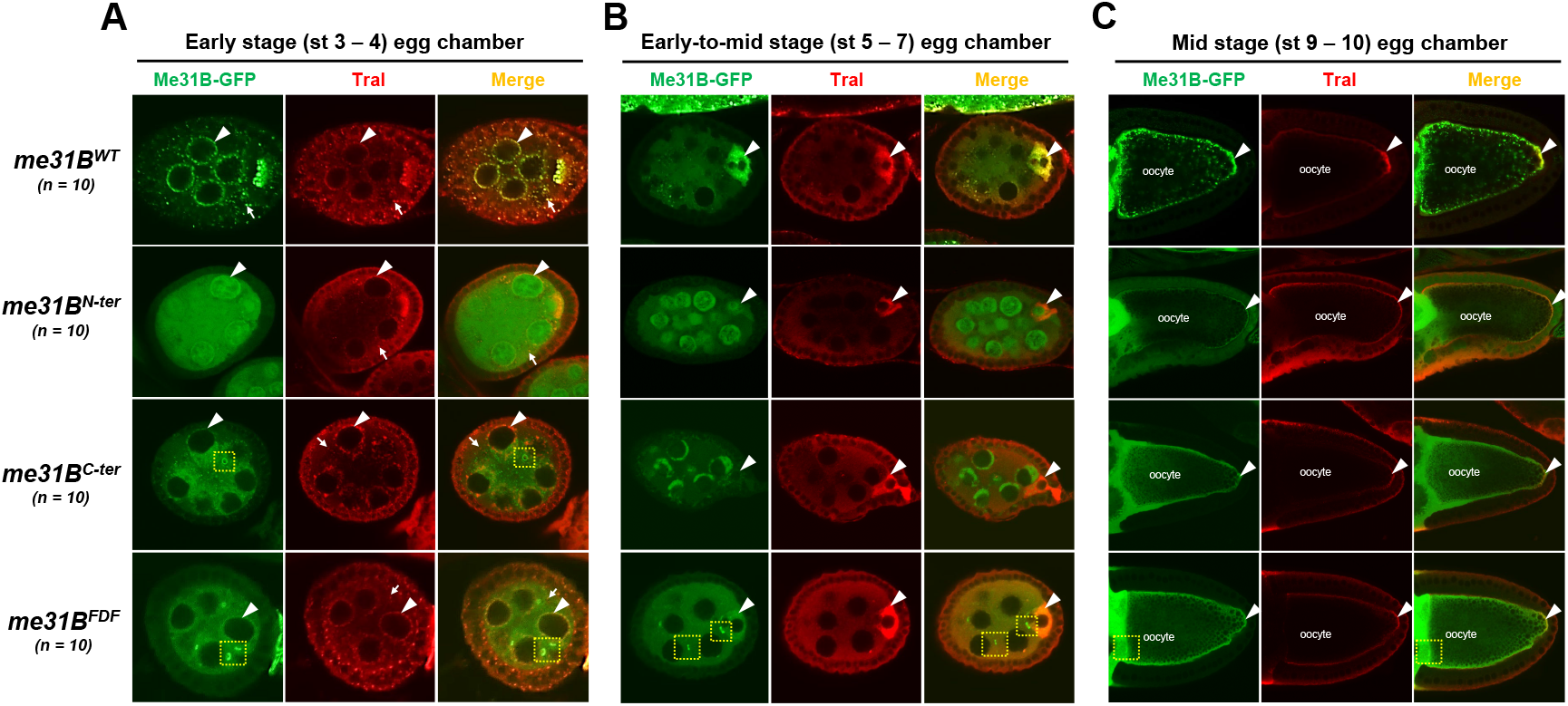
Mutant Me31B proteins in *me31B^N-ter^, me31B^C-ter^, me31B^FDF^* strain show altered aggregation and localization in developing egg chambers. (**A**) In early-stage egg chambers, mutant Me31B-GFP proteins (green channel) in *me31B^N-ter^, me31B^C-ter^*, and *me31B^FDF^* strains are much more diffused in the nurse cell and oocytes, in contrast to the aggregated status of Me31BWT in RNP granules like nuage granules and P-bodies. And none of the three mutant proteins overlap with partner protein Tral. Unlike Me31B mutant proteins, Tral (Red channel) localization are not affected in the three mutants. Particularly, Me31B^N-ter^-GFP proteins are present in the nuclei of nurse cells. Me31B^C-ter^-GFP proteins form fewer numbers and smaller size granules than Me31B^WT^-GFP, and the Me31B^C-ter^ granules do not associate with Tral-marked granules. Nurse cell perinuclear regions (nuage) are indicated by arrowheads. P-body granules marked by Tral are indicated by arrows. Note that Me31B^C-ter^-GFP and Me31B^FDF^-GFP proteins were found in ring-like structures that appear to be ring canals (yellow dashed squares, same for B and C), structures that connect the cytoplasm between nurse cells and oocytes and allow for intracellular transportations. **(B)** In early-to-mid stage egg chambers, mutant Me31B-GFP proteins (green channel) in *me31B^N-ter^* and *me31B^C-ter^* strains do not enrich in the developing oocytes like that in the *me31B^WT^* control. Me31B^FDF^-GFP protein’s enrichment in the developing oocytes is weaker than that in the control. Tral (red channel)’s enrichment in the oocytes is not affected in the three mutants. Developing oocytes are indicated by arrowheads. **(C)** In mid-stage egg chambers, mutant Me31B-GFP proteins of *me31B^N-ter^, me31B^C-ter^, and me31B^FDF^* strains localize to the cortex and the germplasm area at the posterior of oocytes, like the control. However, all three mutant Me31B proteins appear more diffused than the aggregated Me31B^WT^-GFP proteins in the above areas. The germplasm areas are indicated by arrowheads. Tral protein (Red channel)’s localization to the cortex and germplasm was not affected in the three mutants.

About the confounding nuclear localization of Me31B^N-ter^ proteins, we speculate that Me31B may be a nucleocytoplasmic shuttling protein with a Nuclear Localization Signal (NLS) sequence in the N-terminal motif. This NLS sequence leads Me31B into the nucleus, and then the C-terminal motif mediates the protein’s export and retaining in the cytoplasm. This speculation is conceptually supported by the nucleocytoplasmic shuttling activities of DDX6 (human homolog of Me31B) [59] and Xp54 (*Xenopus* homolog) [11] in cell culture models. However, our efforts to identify a Nuclear Localization Signal (NLS) sequence by using prediction tools (NLStradamus [63], cNLS Mapper [64], and SeqNLS Prediction Server [65]) were unsuccessful. Although we were able to locate amino acid sequences in Me31B (KSKLKLPPKDNRFK and CIPVLEQIDP) that are homologous to the putative NLS and Nuclear Export Signals (NES) sequences in DDX6 [59] respectively, experimental evidence is still needed to validate their *in vivo* functionality. We do not exclude the possibility that the N-terminal NLS is just a non-functional sequence that is normally masked in folded, full-length Me31B protein, and the sequence was accidentally exposed to nucleus transportation machinery upon the deletion of the C-terminal motif.

In the early-to-mid stage egg chambers (Figure 6B), we observed that Me31B enrichment in developing oocytes is abolished in the three mutants. In the *me31B^WT^* control egg chambers, Me31B^WT^-GFP and Tral both were highly enriched in the developing oocytes, with the two proteins extensively overlapping (Figure 6B, first row). However, in the *me31B^N-ter^* and *me31B^C-ter^* egg chambers, the mutant Me31B proteins show no obvious enrichment in the future oocytes (Figure 6B, second and third row), while Tral’s enrichment pattern remain unaffected. In the *me31B^FDF^* mutants, the Me31B^FDF^-GFP proteins still showed enrichment in the oocytes, but it is much weaker than the Me31B^WT^-GFP control. We conclude that the Me31B N-ter motif, C-ter motif, and the FDF motif-binding motif are needed for the protein’s proper accumulation into future oocytes at early-to-mid stages. For similar reasons discussed before, we suggest that Me31B’s transport/accumulation into the oocytes depends on its RNA-interaction and protein-binding abilities. This speculation is in line with the observation that Me31B^FDF^ proteins maintained a reduced enrichment in the oocytes: the protein’s intact RNA-interacting motifs or other protein-binding motifs could have enabled an ineffective but functional transportation into the oocytes.

In the mid-stage egg chambers (Figure 6C), Me31B^WT^ proteins were found along the cortex and localized to the germ plasm area of the oocytes (Figure 6C, first row), similar to the Me31B proteins in wildtype strains as previously reported [28]. In the *me31B^N-ter^, me31B^C-ter^, me31B^FDF^* mutants, the three mutant proteins localized to the above areas, but the localized proteins appeared to have less granularity (Figure 6C, bottom three rows), consistent with the diffused status of the mutant proteins in earlier stages.

To further validate that the above localization phenotypes of the mutant Me31B proteins were not a result of defective germline RNP formation, we performed immunostaining against another germline RNP marker, Cup, a protein that complexes with both Me31B and Tral in germline RNPs such as nuage granules, P-body granules, and germplasm granules [28]. We found that, like Tral, Cup’s aggregation and localization into those RNPs remain unchanged in the *me31B* mutants (data not shown).

## Summary

All in all, this study took a mutagenesis approach to investigate the *in vivo* functions of six key motifs of DEAD-box RNA helicase Me31B, and the results are summarized in Figure 7. The three ATPase/helicase motifs (DEAD-box, DVLARAK, and HRIGR) are conserved among the members of the Me31B protein family, and mutations in each of them result in alleles (*me31B^E208A^, me31B^DVLAAAA^,* and *me31B^R385Q^*) that cause dominant female sterility. Their sterility exhibits oogenesis defects like nurse cell dumping defects or egg chamber degeneration and early embryogenesis arrest accompanied by embryo morphological abnormalities. In our attempt to find the molecular-level mechanism by which the ATPase/helicase motif mutations cause sterility, we analyzed the *me31B^DVLAAAA^* heterozygous mutant and showed that the mutation altered *nos* mRNA localization in the germ plasm and decreased *nos* and *osk* transcript levels, highlighting Me31B ATPase/helicase’s function in maintaining germline mRNA localization and transcript level. Our additional analysis of Me31B’s N-ter motif, C-ter motif, and FDF motif-binding motif demonstrated their impotence in fertility, maintaining Me31B protein level and aggregation status, and subcellular localization. The results from this study provided insight on the molecular mechanisms of key functional motifs in *Drosophila* Me31B. Considering the conserved nature of Me31B and its homologs, these data aid in paving the road to understanding the functions and important regions of Me31B family proteins in various cell types and across different species.

**Figure 7.**
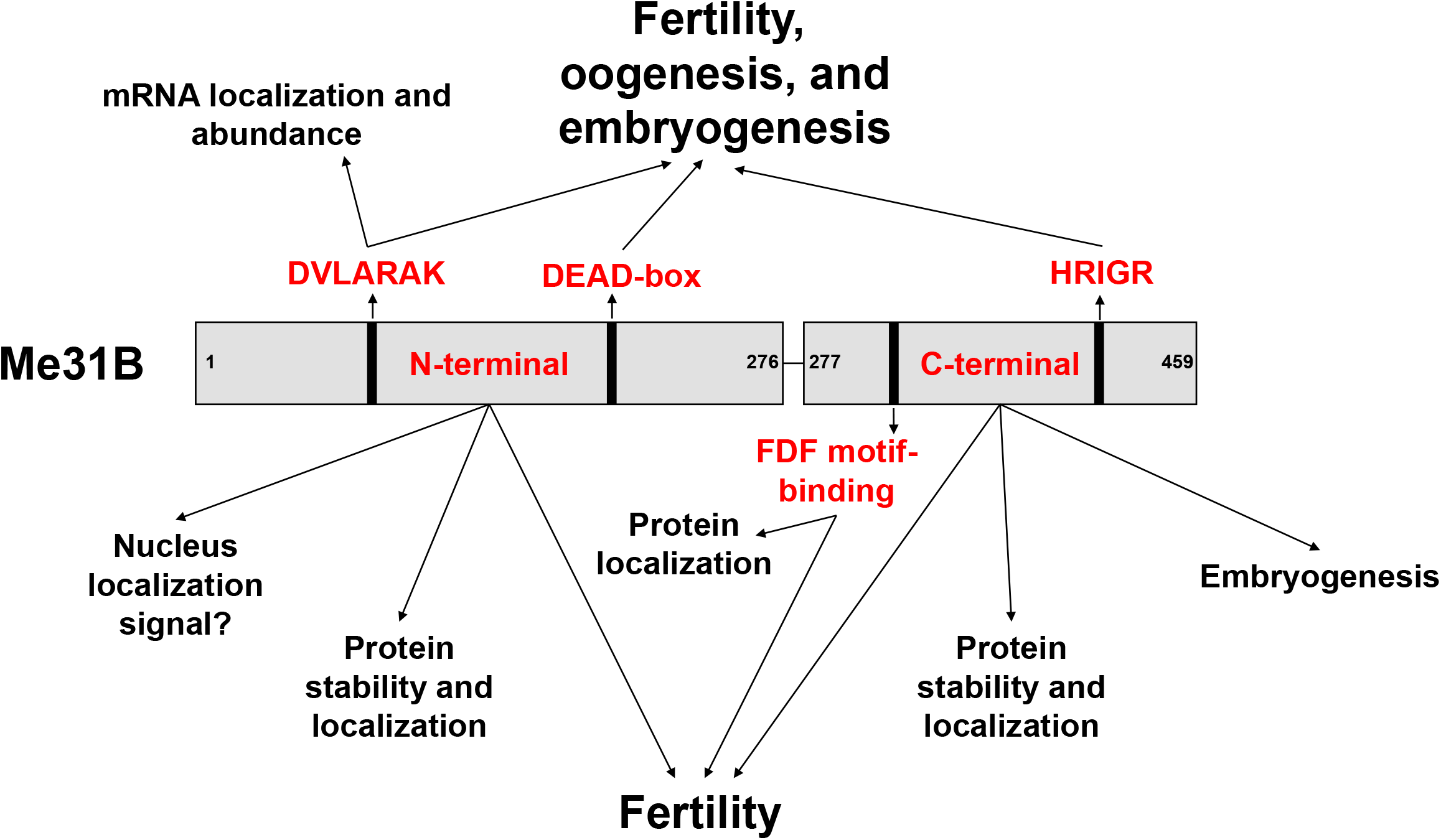
Summary of the Me31B motif functions. This study uses a target-motif-mutation approach to investigate six functionally important domains/motifs of *Drosophila* Me31B. Our characterization of the generated *me31B* mutants revealed ATPase/helicase motifs’ function in female fertility, oogenesis, and embryogenesis (DVLARARK, DEAD-box, and HRIGR motifs). An in-depth analysis of the DVLARAK motif mutation uncovered its function in maintaining *nos* mRNA localization and the transcript level of *nos* and *osk* mRNA levels. We further showed the Me31B N-terminal motif, C-terminal motif, and FDF motif-binding motif’s function in female fertility and their different roles in maintaining Me31B protein level and subcellular localization.

## Materials and Methods

### Fly strain generation by CRISPR gene editing

Mutant *me31B Drosophila* strains were generated by using the previously reported CRISPR procedure [66, 67]. Specifically, CRISPR Optimal Target Finder [66] and DRSC/TRiP Functional Genomics Resources (Harvard Medical School) were used to find gRNA cutting sites flanking the *me31B* gene in the *Drosophila* genome. Then, the found gRNA sequences were cloned into gRNA-expressing plasmid vector pCFD5 (Addgene) according to the suggested protocols. HDR donor plasmids carrying wild type *me31B* gene were constructed by cloning *me31B* gene DNA into pHD-sfGFP-ScarlessDsRed cloning vector (Addgene) according to suggested protocols. In the constructed HDR plasmids, the super-fold GFP (*sfGFP*) gene is positioned in-frame and downstream of the *me31B* gene so that the sfGFP protein is tagged to the C-terminal end of the expressed Me31B proteins. The *DsRed* marker gene is positioned in the intergenic region downstream of the *me31B* gene. HDR plasmid with mutant *me31B* genes was generated by mutating the *me31B* wild type gene in the wildtype HDR donor plasmid by using the Site-Directed Mutagenesis kit (New England Biolabs) according to the manufacturer’s recommended protocols. The resulting HDR plasmids containing different *me31B* alleles are named after the carried mutations (Figure 1). The gRNA-expressing plasmid and the HDR donor plasmids were co-injected into Cas9-expressing strain (Genetivision) to generate desired wild type and *mutant* me31B strains, which were crossed with a 2^nd^ chromosome balancer to establish balanced stocks when possible. The plasmid vectors constructed and the obtained *me31B* strains were validated by sequencing.

### Genetic crosses

Balanced *me31B* wild type and mutant strains were self-crossed to obtain homozygous mutant strains. For the dominant female sterile *me31B* strains, males carrying the mutant allele were crossed with *w^1118^* (Bloomington Drosophila Stock Center 3605) females to obtain heterozygous strains.

### Immunohistochemistry

*Drosophila* ovary immunostaining was performed as previously described [24, 68, 69]. The following antibody dilutions were used: Rabbit-anti-Tral (1:1,000). Donkey-anti-Rabbit-Cy3 secondary antibodies (Jackson ImmunoResearch) were used at 1:500. Images were captured by an Olympus FV3000 confocal laser scanning microscope.

### Western blots

Western blot antibodies were used at the following dilutions: rabbit-anti-GFP (1:100,000), and mouse-anti-α-Tubulin (1:100,000). Secondary antibodies were used at the following dilutions: mouse-anti-rabbit HRP (Jackson ImmunoResearch) (1:10,000 for rabbit-anti-GFP), goat-anti-mouse HRP (Santa Cruz Biotechnology) (1: 50,000 for mouse-anti-α-Tubulin primary antibody). The protein band quantification analysis was performed by using ImageJ (https://imagej.nih.gov/ij/)

### Fertility assay (for *me31B^WT^, me31B^N-ter^, me31B^C-ter^,* and *me31B^FDF^* strains)

Fertility assays were performed according to previously established protocols [60]. Briefly, virgin females from *me31B^WT^, me31B^N-ter^, me31B^C-ter^,* and *me31B^FDF^* strains were collected and allowed to age for 3 – 4 days separately in fly food vials. Each female was then put in a vial in with a *w^1118^* male. After 24 hours, the males were removed from the vials, the fertilized females were transferred to a new vial every day for the next 10 days. The eggs laid and the progenies hatched from each vial were counted. The hatch rate of each vial is calculated by dividing the progenies hatched by the number of eggs laid. An equal amount (15 ml) of fly food medium (Molasses Formulation, Genesee Scientific) was used in each vial.

### Number of egg laying, embryo morphology, and embryo hatchability (*me31B^E208A^, me31B^DVLAAAA^,* and *me31B^R385Q^* strains)

To record the number of eggs laid from *me31B^DVLAAAA^/+* strain, 40 females from the strain were put into a small embryo collection cage with grape-agar plates (Genetivision) in the presence of 10 *w^1118^* males. After letting the flies adjust for 24 hours, the flies were given a fresh grape agar plate, and the eggs laid in the next 24 hours were counted. Control experiments were conducted with *w^1118^* females, and six independent replicates were performed. To observe the morphology and the hatchability of the embryos from *me31B^E208A^/+, me31B^DVLAAAA^/+, me31B^R385Q^/+,* and *w^1118^* control strains, approximately 80 females from each strain were put into a small embryo collection cage in the presence of 20 *w^1118^* males, and the embryos were collected on grape agar plates at 25°C after 24 hours. The embryos were counted and analyzed under a dissection microscope for their morphology. The embryos’ hatchability was calculated 72 hours after they were laid by counting the number of those that developed into larvae (or later stages) or by counting those that failed to develop and then subtracting the failed ones from the total number of eggs laid.

### RNA extraction, cDNA synthesis, and quantitative RT-PCR

Ovarian total RNA was extracted from 5 μl freshly dissected fly ovaries by using Qiagen RNeasy Purification Kit (Qiagen) according to the manufacturer’s instructions. The obtained RNA samples’ concentrations were measured by using NanoDrop 2000c. The RNAs were reversely transcribed to cDNAs by using the High-Capacity cDNA Reverse Transcription Kit (ThermoFisher) according to the manufacturer’s instructions. The synthesized cDNAs were then used for quantitative PCR by using Luna Universal qPCR Master Mix (New England Biolabs). The following PCR Primers were used in this study: *nos* forward 5’ GTCACCAGCAAACGGACGAGATT −3’, *nos* reverse CGGAGCACTCCCGTAGGACAT, *osk* forward 5’- TTGCTGAGCCACGCCCAGAA −3’, *osk* reverse 5’- ACATTGGGAATGGTCAGCAGGAAATC −3’, *rp49* forward 5’- GCTAAGCTGTCGCACAAA, *rp49* reverse 5’- TCCGGTGGGCAGCATGTG −3’. *rp49* RNA was used as the reference. Data analysis was conducted by using the CFX Manager Software (BioRad) and Microsoft Excel.

### smFISH, image analysis, and computational modeling

smFISH was carried out as previously described using published *nos* probe sequences [57, 58, 70]. In summary, ovaries were dissected from yeast-fed females in cold PBS in under 10 minutes and tissues were fixed for 30 minutes in 4% paraformaldehyde and PBS solution. Tissue was incubated with smFISH probes overnight at 37 °C in the hybridization solution previously described [57]. For imaging, egg chambers were mounted in Prolong Glass (Life Technologies) and were allowed to cure for 72 hours at room temperature [57, 58]. A Leica STELLARIS 5 confocal microscope was used for imaging *nos* smFISH experiments that are describe in detail [58]. The identification and quantification of unlocalized single *nos* transcripts and localized *nos* homotypic clusters were carried out using a custom MATLAB (Mathworks) program that has been previously described and published [57, 58]. Confocal images shown in Figure 3 are maximum projections that were filtered by a balanced circular difference-of-Gaussian with a center radius size of 1.2 pixels and surround size of 2.2 pixels as previously done for other germ plasm studies [56, 58]. The total number of homotypic clusters identified are reported in the figure legends. For modeling experiments, we employed a previously published model that simulates the formation of germ granules, including *nos* homotypic clusters [58]. The only modeling parameters that were adjusted were 1) carrying capacity, which is regulated by Osk levels, was set from 1 (wild type) to 0.56, and 2) the pool of *nos* transcript expression was set from 1 (wild type) to 0.61 to mimic the RT-PCR levels reported in our results section [58].

### Statistical Analysis

Reported p-values between average *nos* homotypic cluster sizes were performed using ANOVA test with a post-hoc Tukey test that was calculated using R statistical programming and R Studio using the function aov and TukeyHSD functions [71, 72]. Violin plots were created using the ggplot and ggstatsplot packages [73, 74].

## Acknowledgments

We thank the members of the Gao Lab for discussing and revising this manuscript. We thank Dr. Olivia Rissland and Dr. Akira Nakamura for their kind gifts of antibodies. We thank the Center for Biological Imaging at Kean University for assisting with image acquisition and the members and the Niepielko Lab for their helpful comments and fruitful discussions. Research reported in this publication was supported by the Eunice Kennedy Shriver National Institute of Child Health & Human Development of the National Institutes of Health under award no. R15HD102960 to MGN and 1R15HD092925-01A1 to MG.

## Author Contributions

MG conceived and directed the project. MG and MGN wrote the manuscript with input from all authors. EK, MP, IF, and NM performed the fertility assays. AM, CD, and JE generated the CRISPR strains. MGN, DAD, and BMO performed the microscopic analysis, smFISH experiments, and modeling. KS and YM performed the confocal microscopy experiments. BP, ARM, and DG performed the Western blots and data analysis. All authors discussed the results and commented on the manuscript.

## Data Availability

All the data presented in this study are included in this manuscript or its supplementary information.

