## Supplementary Figure for "An *In Vivo* Analysis of the Functional Motifs of DEAD-box RNA Helicase Me31B in *Drosophila* Fertility and Germline Development"

### Supplementary Figure 1: *me31B$^{DVLAAAA}$*/+ strain showed decreased egg lay

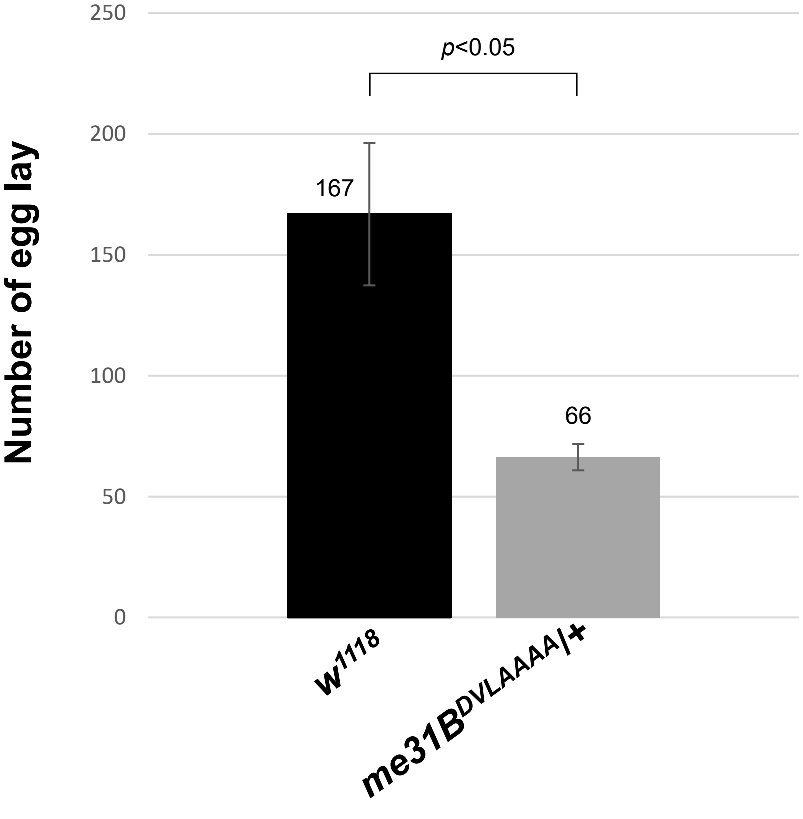


**In this assay condition, *me31B$^{DVLAAAA}$*/+ strain lay an average of 66 eggs, which is 40% of that from the *w$^{1118}$* control strain ($p < 0.05$)**

#### Supplementary Figure 2: Additional and uncropped biological replicate images of main text Figure 5.

**Biological replicate 1: Anti-GFP and anti-α Tubulin Western blots with *me31B$^{WT}$*, *me31B$^{N-ter}$*, *me31B$^{C-ter}$*, *and me31B$^{FDF}$* ovaries.**

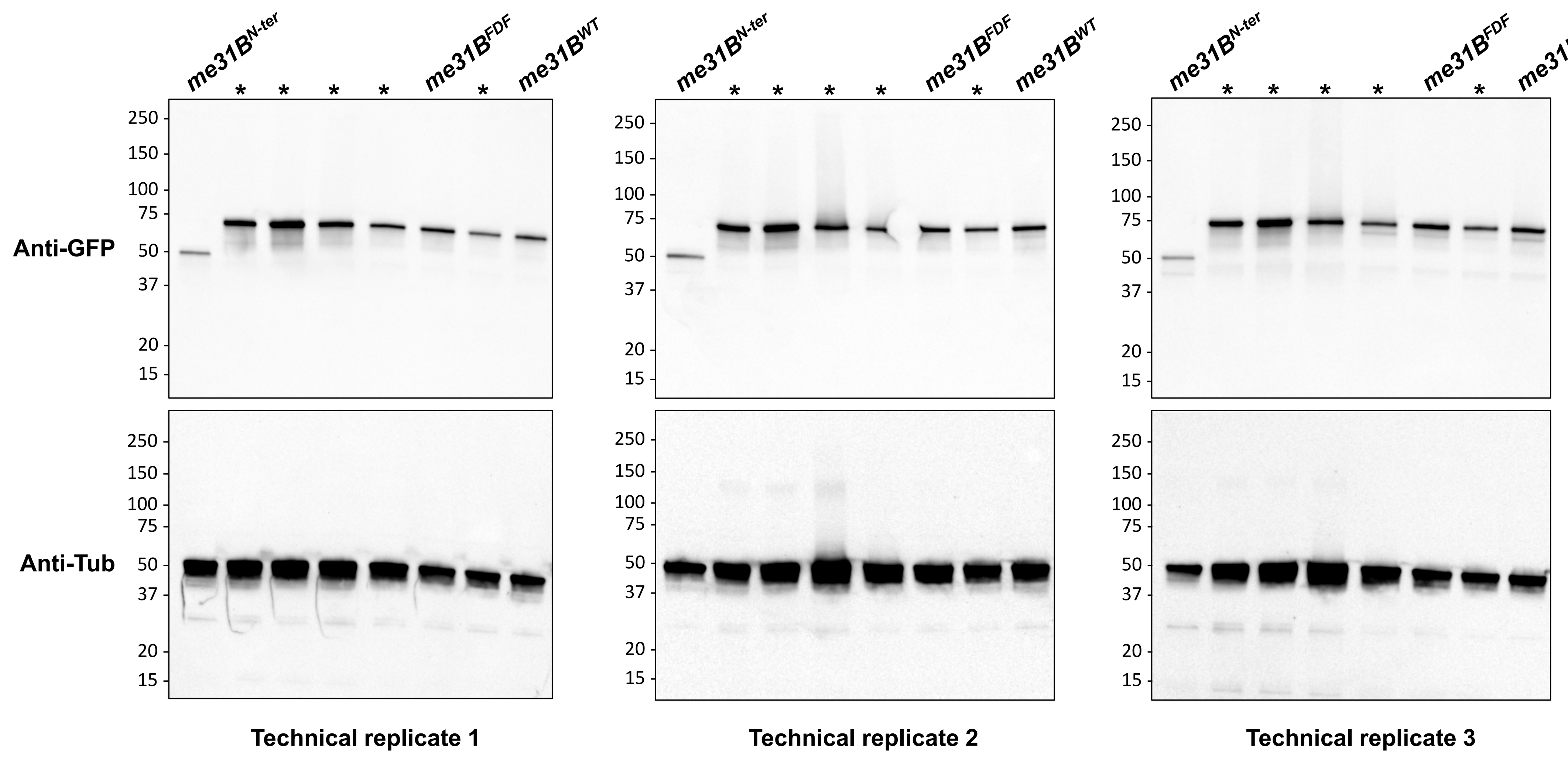


(*Asterisks represent other *me31B* strains not involved in this study. Same below.)

**Biological replicate 2: Anti-GFP and anti-α Tubulin Western blots with *me31B$^{WT}$*, *me31B$^{N-ter}$*, *me31B$^{C-ter}$*, *and me31B$^{FDF}$* ovaries.**

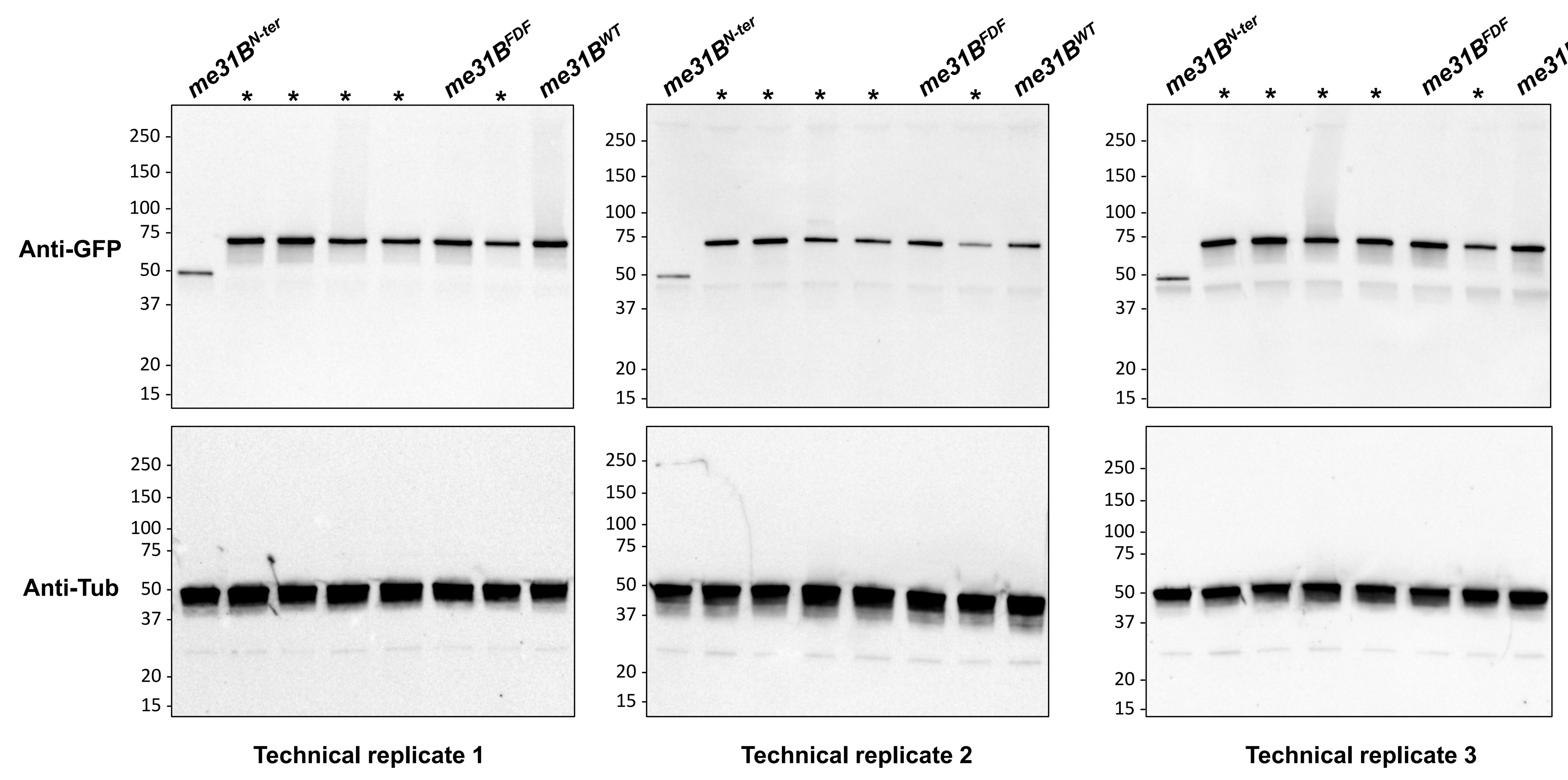


**Biological replicate 3: Anti-GFP and anti-α Tubulin Western blots with *me31B$^{WT}$*, *me31B$^{N-ter}$*, *me31B$^{C-ter}$*, *and me31B$^{FDF}$* ovaries. The cropped images in main text Figure 5 are highlighted by the yellow squares below.**

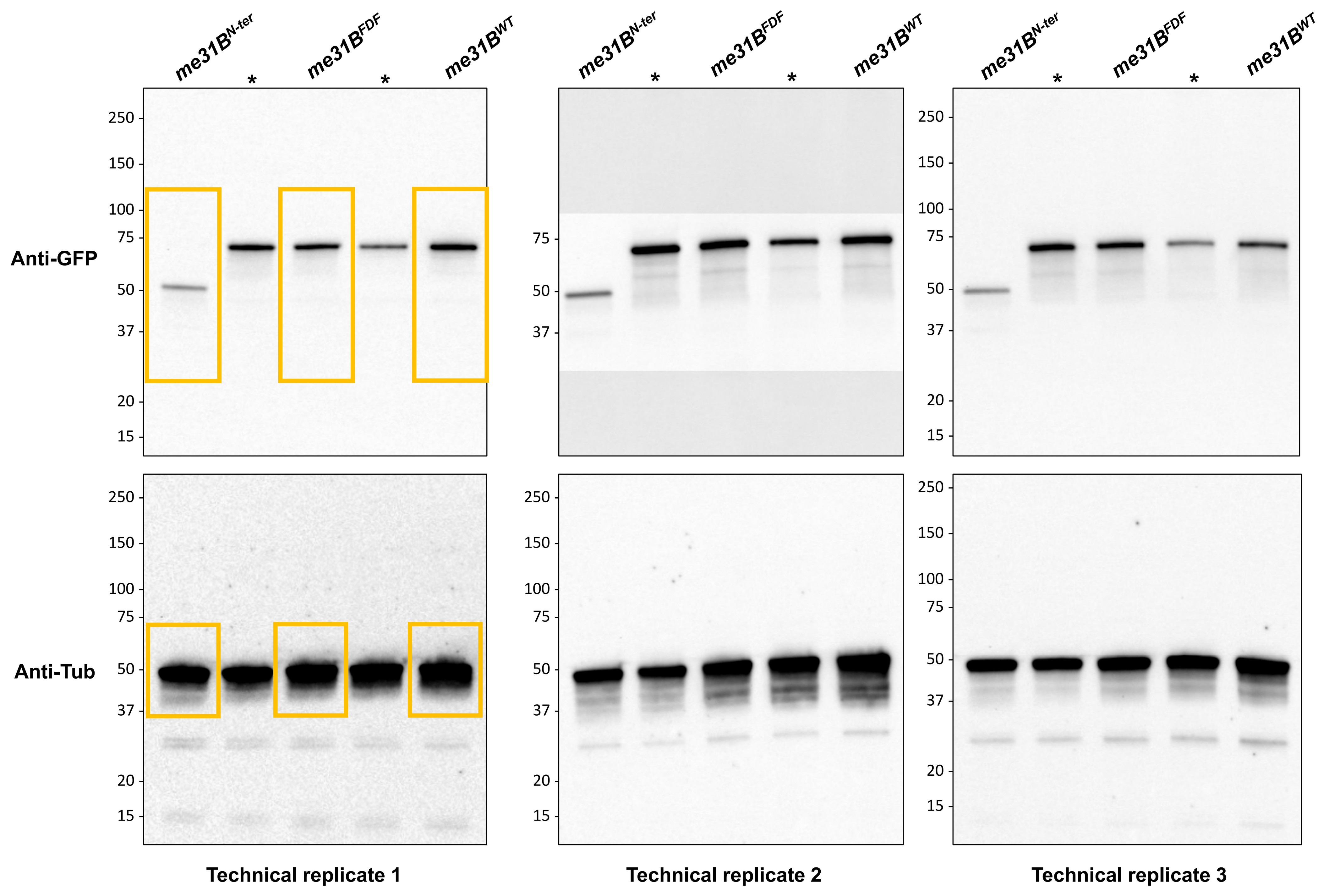


**Biological replicate 1: Anti-GFP and anti-α Tubulin Western blots with *me31B$^{WT}$* and *me31B$^{C-ter}$* ovaries. The cropped images in main text Figure 5 are highlighted by the yellow squares below.**

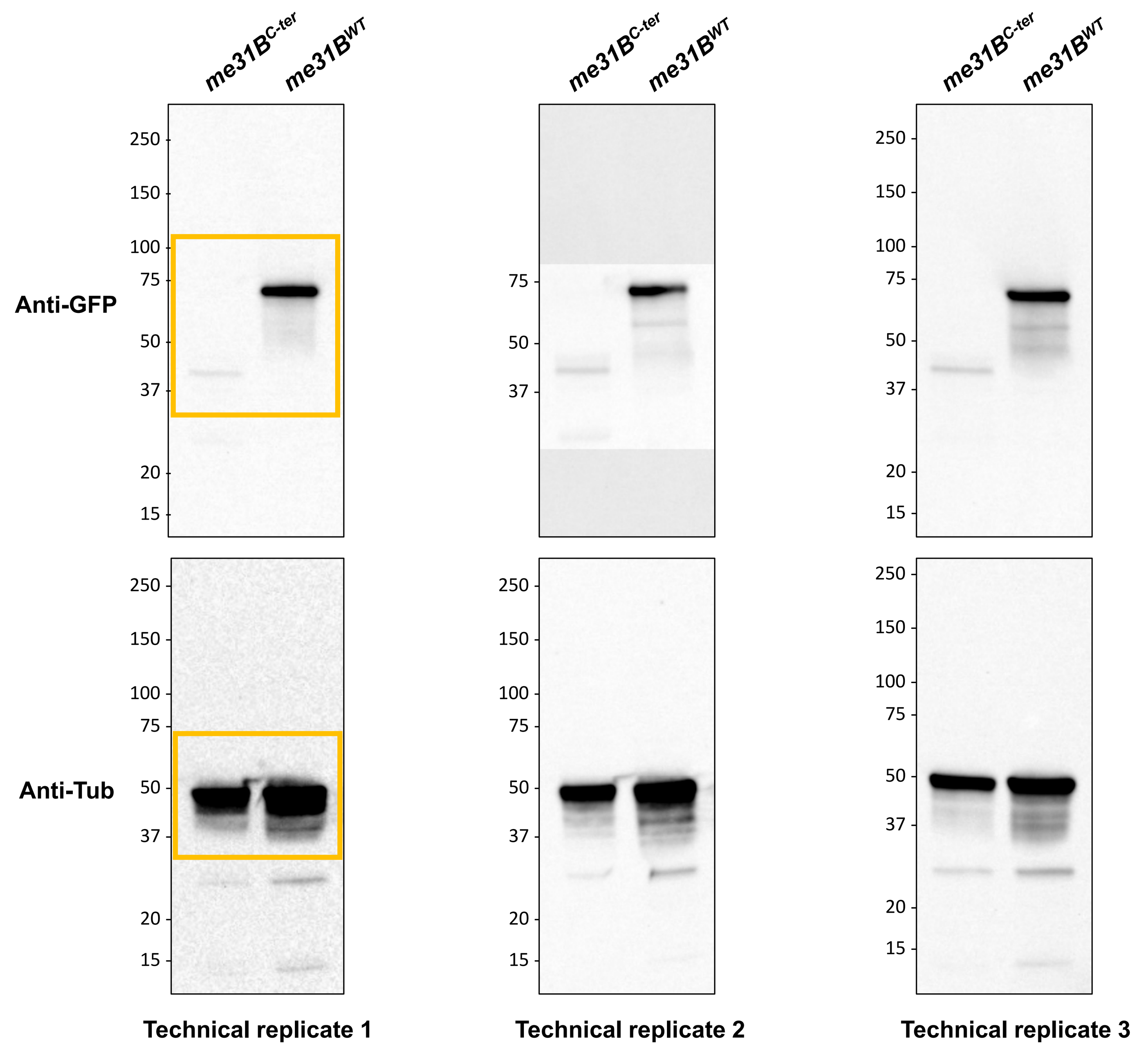


**Biological replicate 2: Anti-GFP and anti-α Tubulin Western blots with *me31B$^{WT}$* and *me31B$^{C-ter}$* ovaries.**

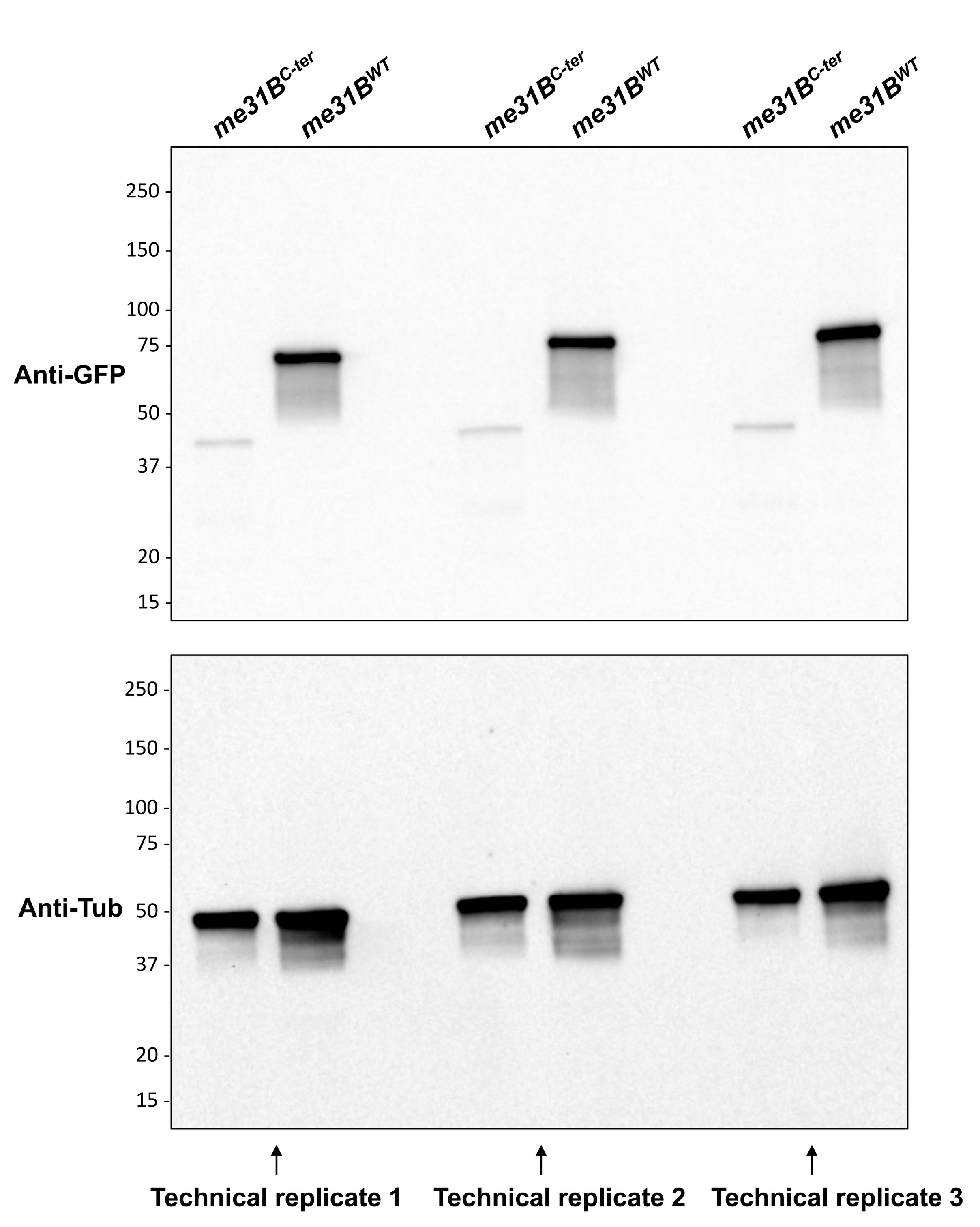


**Biological replicate 3: Anti-GFP and anti-α Tubulin Western blots with *me31B$^{WT}$* and *me31B$^{C-ter}$* ovaries.**

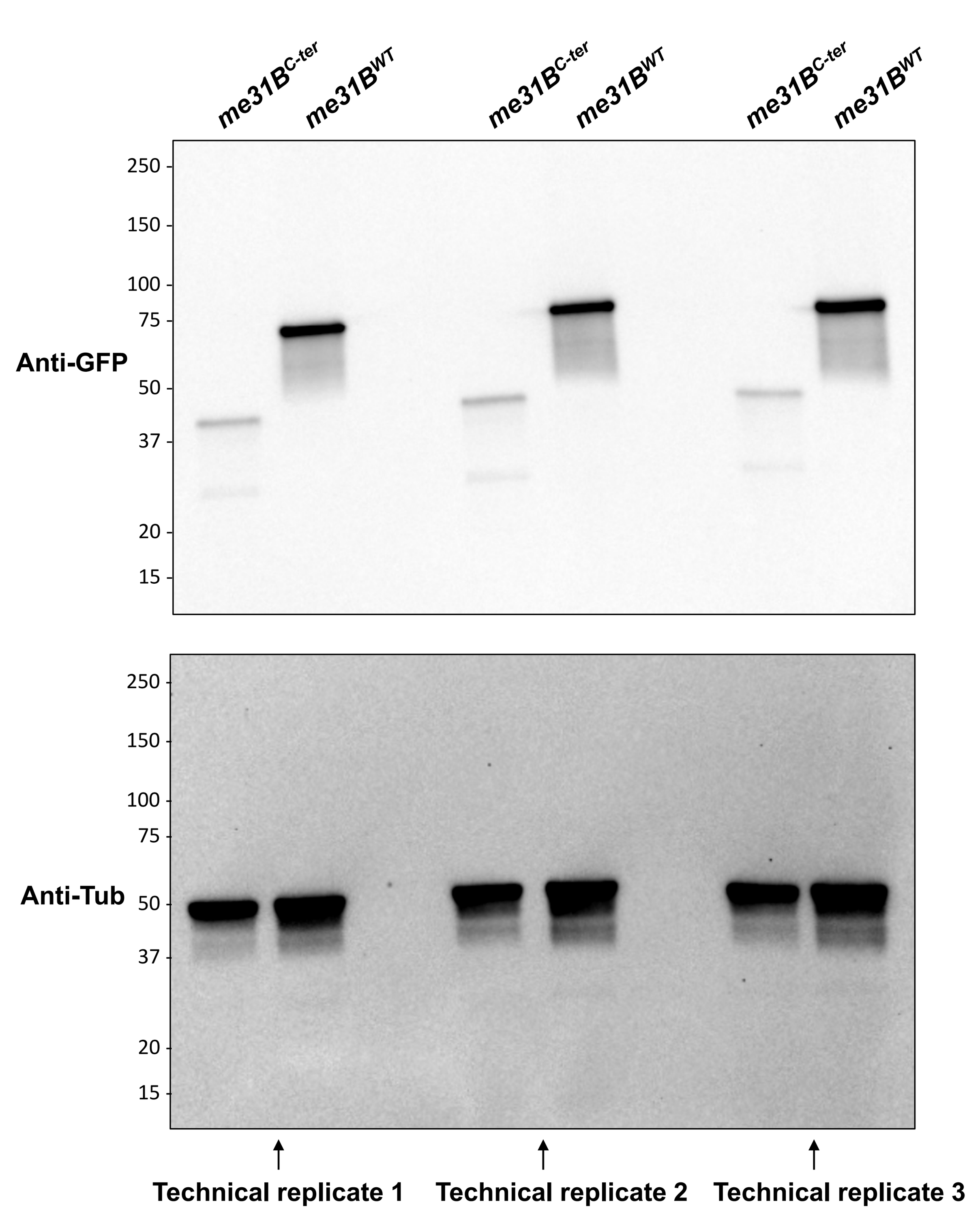

### Supplementary Figure 3. Me31B$^{N\text{-}ter}$-GFP localize to nurse cell nuclei

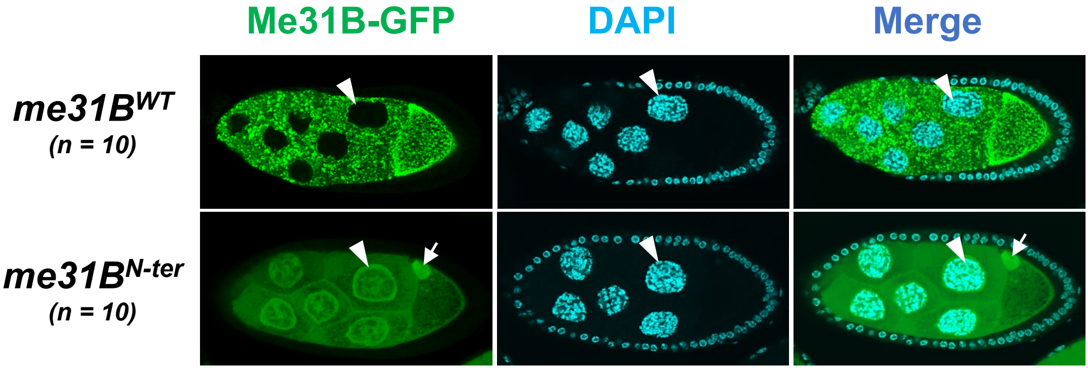


Me31B$^{N\text{-}ter}$-GFP (green channel) proteins are present in the nuclei of nurse cells (indicated by arrow heads) in the *me31B*$^{N\text{-}ter}$ strain, while Me31B$^{WT}$-GFP proteins are found only in the cytoplasm of nurse cells and oocytes. Nucleus DNA are stained by DNA stain DAPI. Nurse cell nuclei are indicated by arrow heads. Interestingly, Me31B$^{N\text{-}ter}$-GFP proteins are also found as a small accumulation in the dorsal-anterior corner of the developing oocyte (indicated by arrows), but this structure does not seem to contain DAPI-stained DNA.
